## Supplementary material for "TGF-β blockade drives a transitional effector phenotype in T cells reversing SIV latency and decreasing SIV reservoirs *in vivo*": Movie_9

### Supplemental Methods

#### ImmunoPET/CT Analysis

Multimodal registration and ROI segmentation were executed using the MIM software (MIM Software, Inc., Cleveland, OH) for PET and CT images, encompassing 250 to 300 fractions, depending on the macaque's body size. PET and CT images were fused to quantify changes in SUV (Standardized Uptake Value) within Regions Of Interest (ROI), with PET SUV scale determined by overall signal intensity and CT scale optimized for tissue visibility. Segmentations were performed for various anatomical regions: Whole Body, Liver, Lymph nodes, Gut, Kidneys, Spleen, and Vertebrae. The external body was outlined for the Whole-Body contour, while the GRADIENT method PET-Edge and interactive contouring tools were employed for other ROIs. Liver segmentation initially utilized PET-Edge in PET coronal-sectional view, followed by interactive delineation for the structure and the corresponding signals. Kidneys, Spleen, and Vertebrae volume were determined based on CT scans in the cross-sectional view, incorporating PET signals into contours. Notably, the analysis for vertebrae was confined to the range from the thoracic vertebrae to the lumbar vertebrae. To quantify the signal in the axillary lymph nodes, the data-point in each macaque featuring the strongest PET signal was chosen as a basis for an in-house Multi-Time-Point Contour Transfer Workflow, followed by interactive contouring of the signal to all time points. The workflow integrated CT scans to establish and map specific ROIs in subsequent scans, thereby accounting for any changes in the animal's orientation at other time points through manual adjustments. To quantify the signal in the gut tissue, the body segment below the stomach to above the cervix was initially isolated. The Gut's SUV was then calculated by extracting the spleen, both kidneys, liver, and bones within the designated body region using Boolean operations. An example of ROIs for each organ and how they were created is shown in Fig S2 and Movies 9 and 10.

#### Clinical Pathology at NIRC

1mL of blood was collected in EDTA tubes and tested for a standard hematology panel (complete blood count - CBC) using a BECKMAN COULTER DxH600 hematology analyzer. An additional 1mL of blood was collected into a SST tube, centrifuged to isolate serum, and analyzed for a comprehensive serum chemistry panel using a Siemens's Dimension Clinical Chemistry Analyzer.

**Cell isolation from tissues, flow cytometry staining and analysis.**

Rectal biopsies were incubated in HBSS containing 2mg/mL Collagenase IV (Worthington Biochemical) and 1mg/ml of Human Serum Albumin (Sigma-Aldrich), shaking at 37°C for 30 minutes. The resulting cell suspension was passed through a 40μm cell strainer and washed with PBS. Cells from lymph nodes were isolated by passing the collected tissues through a 70μm cell strainer. Isolated cells were frozen in liquid nitrogen for phenotyping and stimulation experiments at a later time.

Peripheral blood mononuclear cells (PBMCs) collected from 8 macaques at 7 time points spanning Cycle 1 to Cycle 4 (refer to Table S6) were employed for high-parameter flow cytometry staining. Cells from before and after cycle 1 and at the end of cycle 3 were used for Tfh/Treg assay. In both cases, cell fixation and permeabilization for intracellular staining were carried out using the FoxP3 Staining Buffer set (Thermo Fisher Scientific, Carlsbad, CA). The high-dimensional panel consisted of antibodies outlined in Table S4, catering to requirements for surface and intracellular staining, as well as viability assessment. Each antibody was individually titrated to ensure optimal staining conditions. Data acquisition was performed using the FACSymphony A5.2 Spectral Analyzer. Following data collection, spectral compensation was implemented utilizing FlowJo™ v10.8 Software (BD Life Sciences). 60,000 live singlet lymphocytes from each of samples was downsampled to prevent disproportionate representation by samples with higher cell counts, ensuring balanced subsequent analysis. To mitigate batch effects, we utilized the swiftReg Neighbor-Dependent Cluster Registration (NDCR) method available within the SWIFT package(*45*). This method effectively addresses batch variations while preserving the inherent biological diversity. The NDCR method is particularly adept at registering samples without requiring an internal reference, achieving correction through a calculated value shared among clusters with comparable channel values. Furthermore, we implemented the Peak Extraction and Cleaning Oriented Quality Control (PeacoQC) procedure[73] in FlowJo™ v10.8 to eliminate low-quality events arising from potential issues like clogs, speed fluctuations, and slow sample uptake. FlowSOM FlowJo plugin was employed for automated cell population identification, preceded by the utilization of Phenograph to obtain optimal cluster numbers for dimensional reduction, for the merged data of All Time points and BC1 vs AC4 merged data respectively. The Supplementary Figure S6 A heatmap illustrates the median fluorescence intensity (MFI) values for all markers across all clusters. A tSNE plot, generated in FlowJo™ using the markers employed in the FlowSOM analysis, is presented, with coloring based on the FlowSOM clustering or two specific time points, BC1 vs AC4 in Fig 4G.

Statistical analysis of Spectral flow cytometry data was executed using Python. To discern disparities in the 36 clusters resulting from FlowSOM between BC1 and AC4, a Wilcoxon test with FDR correction was employed. To evaluate variations within the 38 clusters arising from FlowSOM analysis across multiple time points, a repeated measures ANOVA was conducted with Holm-Sidak correction for multiple comparisons.

**Single-cell RNAseq Analysis:**

Cells were isolated from LN biopsies as described above and fixed with the Parse fixation kit, barcoded and sequenced at the NUSeq Core. Libraries were validated by capillary electrophoresis on a Fragment Analyzer (Agilent),and sequenced using 1 S4 flow cell of the Illumina NovaSeq 6000 yielding >20,000 reads/cell. Raw sequencing data were demultiplexed, aligned to Mmul10 and QC with the Parse pipeline. The Partek software was used for the remaining QC. We filtered out cells with less than 400 and more than 8000 features, excluded features with 0 reads in >99.99% of cells, scran deconvolution was used for normalization and cell classification was based on gene expression. Since mitochondrial counts were already below 2% after these filters, we did not filter further. Scran deconvolution was used for normalization and principal component analysis (PCA) and Uniform Manifold Approximation and Projection (UMAP) for data dimensionality reductions and visualization. Cells were classified as T cells if expressed any CD3 (CD3D, CD3E, or CD3G), but no CD19, PAX5 or MS4A1; B cells if they expressed CD19 or MS4A1 or CD79A or PAX5 but no CD3; NK cells if they expressed NCAM1 or NKG7 or KLRB1 but no CD3, CD14, MS4A1, CD19 or CD68. Finally, macrophages were CD68+, but CD19- CD79A- LAMP3- and CD3-. In a separate classification, CD4 T cells were classified as expressing CD3 and CD4 or CD8 (CD8A or CD8B), but no CD19, MS4A1 or CD68. Finally, a third classification classified GC B cells as A4GALT (CD77)+ or BCL6+ or RGS13+ and CD19+ or PAX5+ or MS4A1+, but CD3-; naïve B cells as TCL1A+ or CCR7+ and CD19+ or PAX5+ or MS4A1+ and CD3-; Tfh cells as expressing CD3 and BCL6+ or CXCR5+ AND CD19-, PAX5-, MS4A1-. Hurdle models were used to compare feature counts in each cell subset before and after treatment and obtain a DEG list. Gene set enrichment with the Partek tool was performed on all significant (FDR<0.05) DEG.

An alternative analysis based on the R package Seurat (version 4.3.0.1) (http://satijalab.org/seurat/) for single-cell expression analysis, as also performed to ensure the accuracy of the Partek analysis and conclusions. The reads for each sample were loaded using ReadParseBio function and identified Rhesus macaque mitochondrial reads using the scCustomize package (version 1.1.1) (https://doi.org/10.5281/zenodo.5706430 ). We filtered out cells with ≤200 genes and for mitochondrial gene fraction dependent on the sample. We normalized all samples using SCTransform(*75*). For downstream analysis, we focused on T cells, defined as expressing CD3 and not CD19 nor PAX5 nor MS4A1. We performed integration according to the Satija lab's integration workflow (https://satijalab.org/seurat/articles/integration_introduction.html). Once the datasets were integrated, we generated data dimensionality reductions by PCA and UMAP, using the first 35 principal components for UMAP generation and a cluster size of 0.5. Next, we further subset the data to isolate CD4$^+$ T cells, defined as T cells expressing CD4, and separately CD8$^+$ T cells, defined as T cells expressing either CD8A or CD8B. We calculated the differentially expressed genes comparing W37 to W35 using Wilcoxon matched pairs test. Together, we observed broad agreement with the independent analysis performed with Partek described above, although there were some differences. In CD4$^+$ T cells, we found 2,022 DEGs with a $\log_2$FC of 0.5 (Adj P-value <0.05) while 2,043 DEGs were in CD8$^+$ T cells and 2,339 DEG were in T cells, with 172 downregulated genes in the CD8$^+$ T cells as compared to 187 in the CD4$^+$ T cells and 74 in T cells. These DEGs covered 36% for T cells and 33% for CD4$^+$ T cells from the Partek analysis. Amongst the AP-1 family genes, the log2FC for this analysis vs Partek analysis was as follows: JUNB (T cell: -1.06 vs 0.06), JUN (T cell: -0.22 vs 0.21), FOS (T cell: 1.21 vs 0.16), FOSB (T cell: -0.70 vs 0.25), FOSL2 (T cell: 2.16 vs 0.01), MAF (T cell: 3.70 vs 0.01), BATF (T cell: 4.70 vs 0.01), ATF3 (T cell: 0.60 vs 0.06), MAFF (T cell: 3.97 vs 0.00). Although $\beta$2 macroglobulin and RPL13 were not differentially expressed in our analysis, the fold change was in the same direction as Partek analysis (T cell: log2FC 1.77 and log2FC 0.03 of SCT normalized counts, respectively). While STAT1 was upregulated in CD4$^+$ T cells (log2FC 0.88), it was downregulated in CD8$^+$ T cells (log2FC -0.72). Although TCF7 was not statistically significantly differentially expressed in CD8$^+$ T cells, the W37 vs W35 SCT normalized expression of TCF7 was in the same direction as Partek analysis (log2FC -0.79).

**Barcode sequencing and analysis**

RNA isolation from plasma was performed using a QIAamp Viral RNA mini kit (Qiagen). Complementary DNA (cDNA) was generated with Superscript III reverse transcriptase (Invitrogen) and an SIV-specific reverse primer (Vpr.cDNA3: 5'-CAG GTT GGC CGA TTC TGG AGT GGA TGC-3'). The cDNA was quantified via qRT-PCR using the primers VpxF1 5'-CTA GGG GAA GGA

CAT GGG GCA GG-3' and VprR1 5'-CCA GAA CCT CCA CTA CCC ATT CATC with labeled probe (ACC TCC AGA AAA TGA AGG ACC ACA AAG GG). Prior to sequencing, PCR was performed with VpxF1 and vprR1 primers combined with either the F5 or F7 Illumina adaptors containing unique 8-nucleotide index sequences for multiplexing. PCR was performed using High Fidelity Platinum Taq (ThermoFisher) under the following conditions: 94˚C, 2 min; 40 Å~ (94˚C, 15 s; 60˚C, 1.5 min; 68˚C, 30 s); 68˚C, 5 min. The multiplexed samples and Phi X 174 library were prepared via standard protocols and sequenced on a MiSeq instrument (Illumina).

For viral population analysis, Shannon Entropy of the viral barcodes present in each sample was used to measure the diversity of the viral populations. Shannon Entropy was calculated in R v4.2.2 using the barcode frequencies applying the formula: $Sh = \Sigma\left(-Pi * \log 2\,(Pi)\right)$; where Sh is Shannon Entropy calculated for each sample, and Pi is the frequency of each barcode. To test for changes in Sh before and after Galunisertib treatment, we fitted linear mixed effects models for each compartment and treatment cycle assuming a log-normal distribution of Sh, using before and after treatment time point as predictor and controlling for Sh value at peak viremia, and within animal correlation, the latter as a random effect. Modeling was performed using the lme4 v1.1-34 R package.

### Supplemental Figures Legends

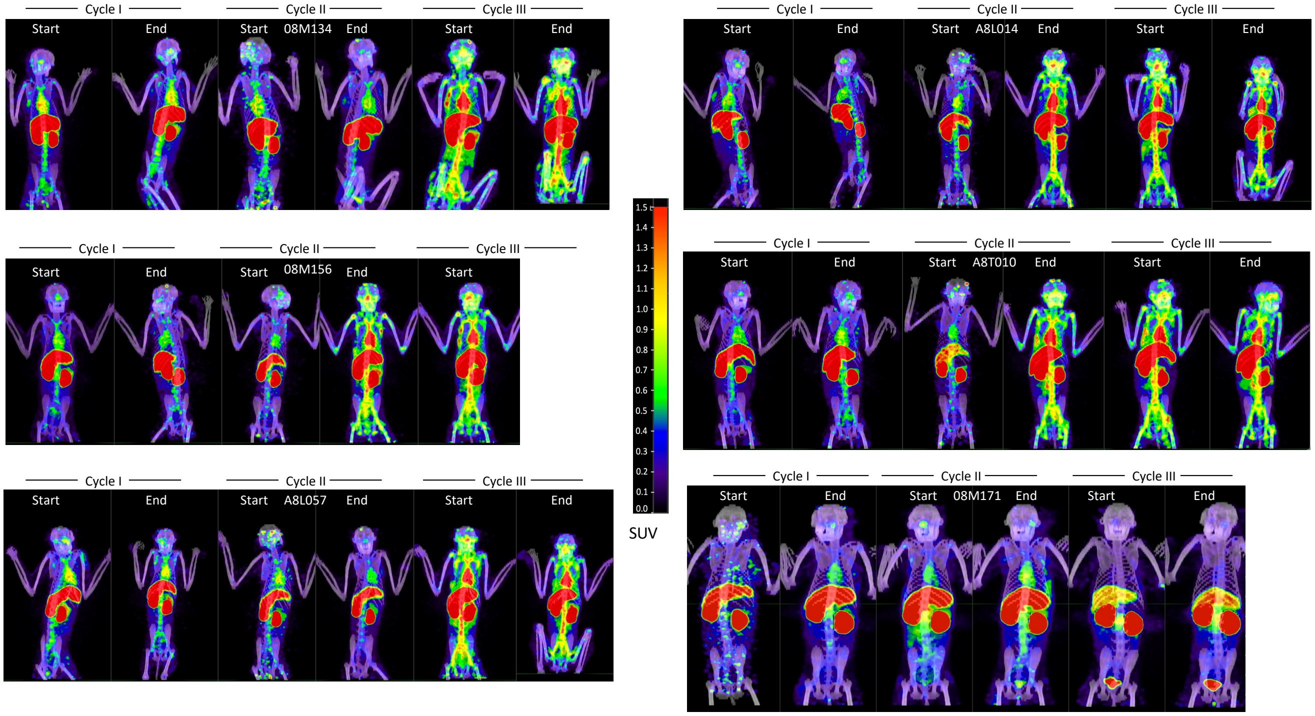


**Figure S1. Representative images of PET/CT scans performed in all macaques before and after each galunisertib cycle.** The $^{64}$Cu-DOTA-Fab$_2$(7D3) probe was injected ~24hrs before PET/CT scan before and at the end of each of the first 3 galunisertib cycles. Representative images from the PET/CT scans are shown for all the macaques not shown in the main Figure 1. The last scan of 08M156 (AC3) did not transfer complete files due to an error on the scanner.

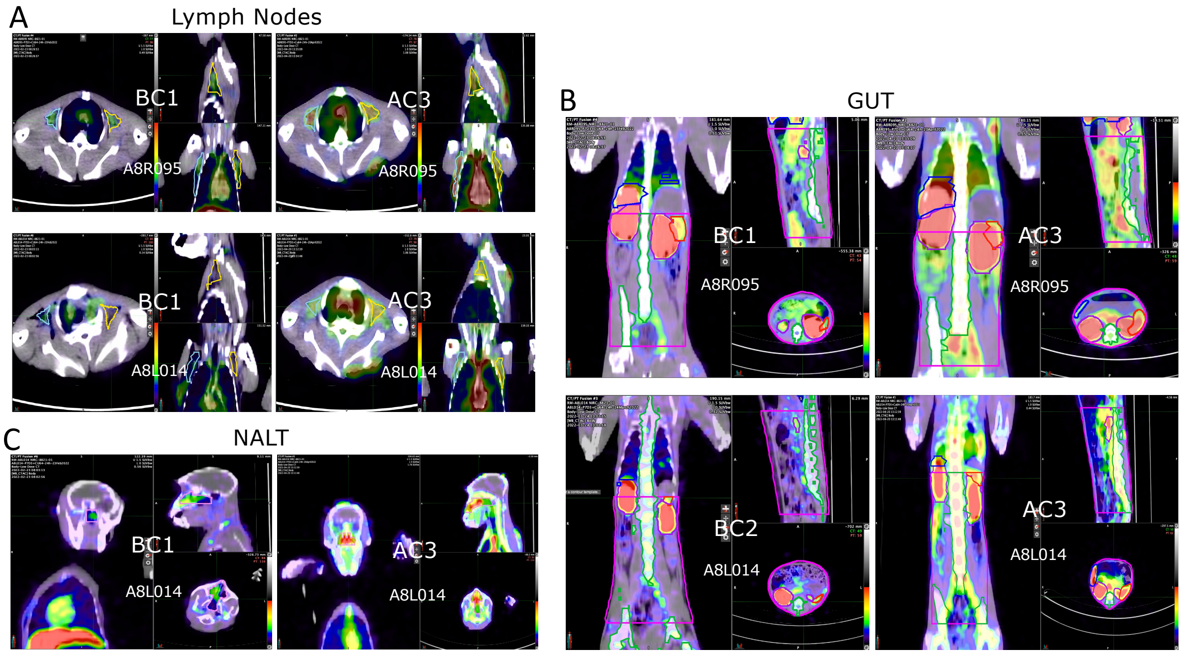


**Figure S2. Examples of ROIs generation for gut, LN and NALT.** A-C) Snapshot representing Maximum Intensity Projections (MPIs) generated using the MIM software and visualized with the color scale PET Rainbow. The MIP were scaled to the same physical size at all time points in all monkeys using the dimensions to the scanner's visible range, and the scanner bed was removed from the CT scan. Examples of ROI for different tissues are shown at 2 time points for 1-2 randomly selected monkey. A) For delineating the axillary lymph nodes ROI (blue line in A), the scan for each animal displaying the most robust signal was contoured on the axial plane based on the Axillary level 1 contour in TCF Van Heijst Phys. Med. Biol 2017, and transferred to other time points using a multi-time-point workflow in MIM. Small adjustments were made to account for CT misalignment. B) The gut region (pink line) was identified from beneath the stomach down to the cervix on the coronal view. To isolate the gut as the region of interest (ROI), specific exclusions were made using Boolean gates to remove the kidneys (purple line), liver (blue line), spleen (red line), and bones (green line) along with their associated signals from within the gut area. All contouring processes were conducted using the pen tool within MIM. C) The delineation of the nasal-associated lymphoid tissue (NALT, pink line) was performed on PT scans, specifically within the nasal cavity extending from the external naris to the nasopharynx. This contouring process involved utilizing the pen tool in MIM across both sagittal and axial views.

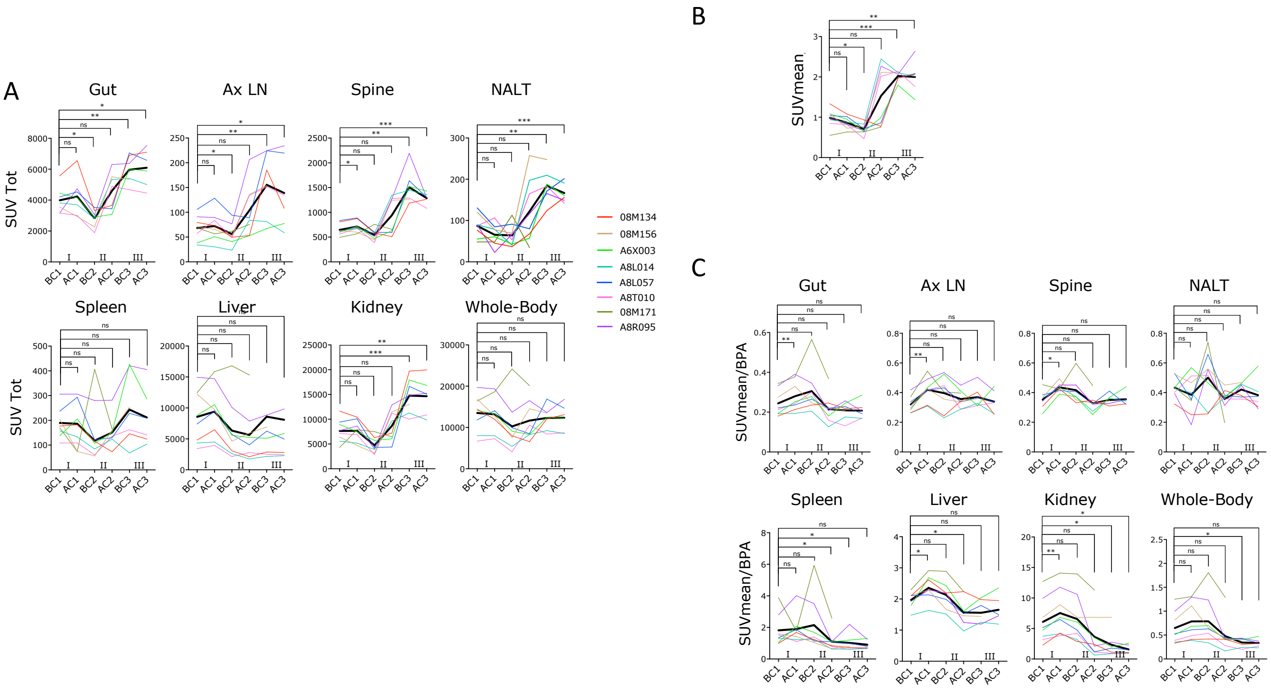


**Figure S3. SUVTotal, blood pool activity (BPA) and SUVmean normalized for BPA.** A) SUV Total is shown for the tissue/organ of interest to demonstrate changes in overall radiotracer uptake in the region. B) Blood pool activity SUVmean (BPA) was measured with a 2D circular region of interest (ROI; 6mm diameter) within the left ventricle cavity using the 3D brush tool available in the MIM software on the chosen CT slice. C) SUVmean for each tissue normalized for BPA are shown. Values were analyzed with mixed-effect analysis. P–values were calculated for comparison of each time point with the before cycle 1 time point (BC1; AC1= after cycle 1, BC2=

before cycle 2; AC2= after cycle 2; BC3= before cycle 3; AC3= after cycle 3; Holm-Sidak multiple comparison correction; *p≤0.05 **p≤0.01 ***p≤0.01)

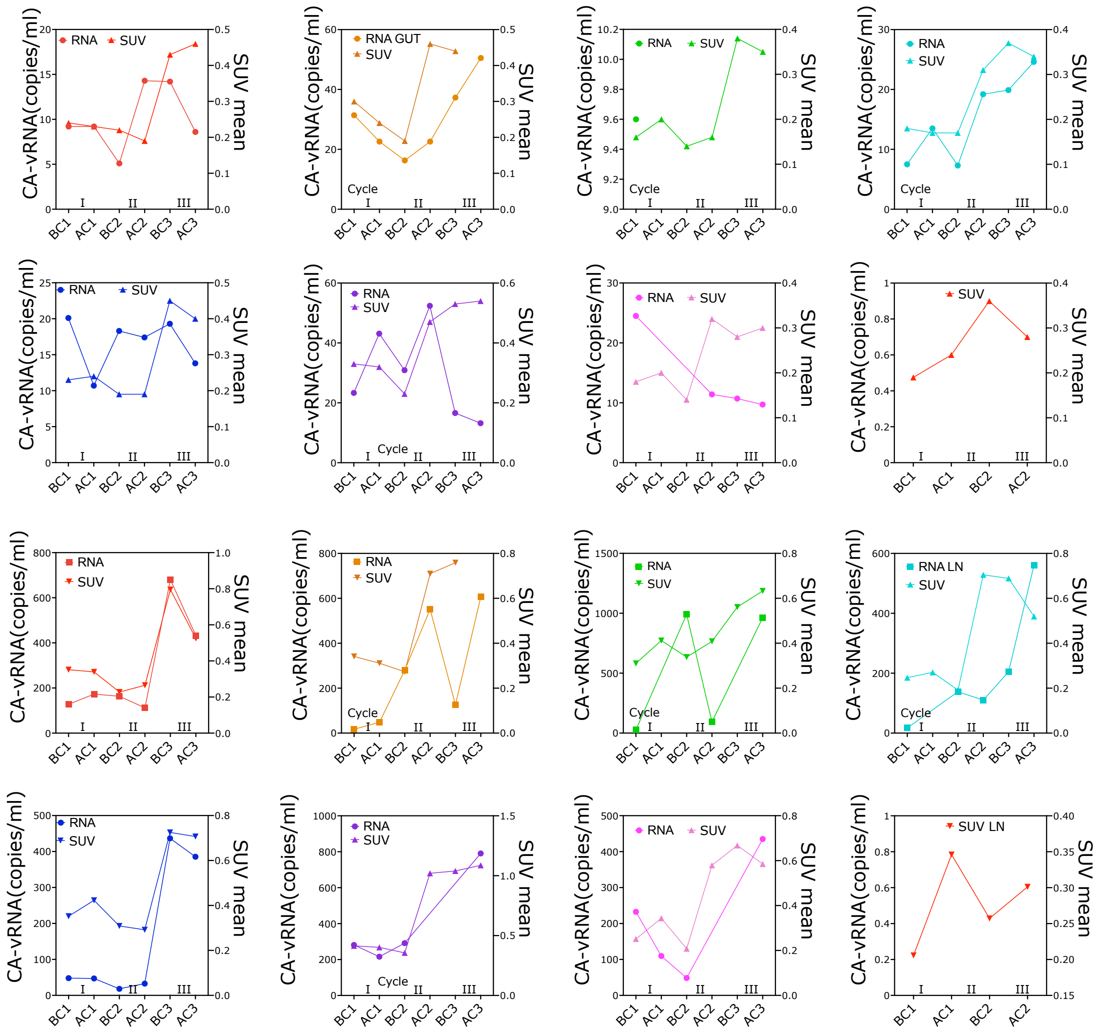


**Figure S4 Relationship between vRNA and SUV in the gut and LN.** SUVmean for the gut and LN ROIs are plotted together with cell-associated vRNA in colorectal biopsies (A) and FNA (B) at the specified time during galunisertib treatment (BC1= before cycle 1, AC1= after cycle 1, BC2= before cycle 2; AC2= after cycle 2; BC3= before cycle 3; AC3= after cycle 3).

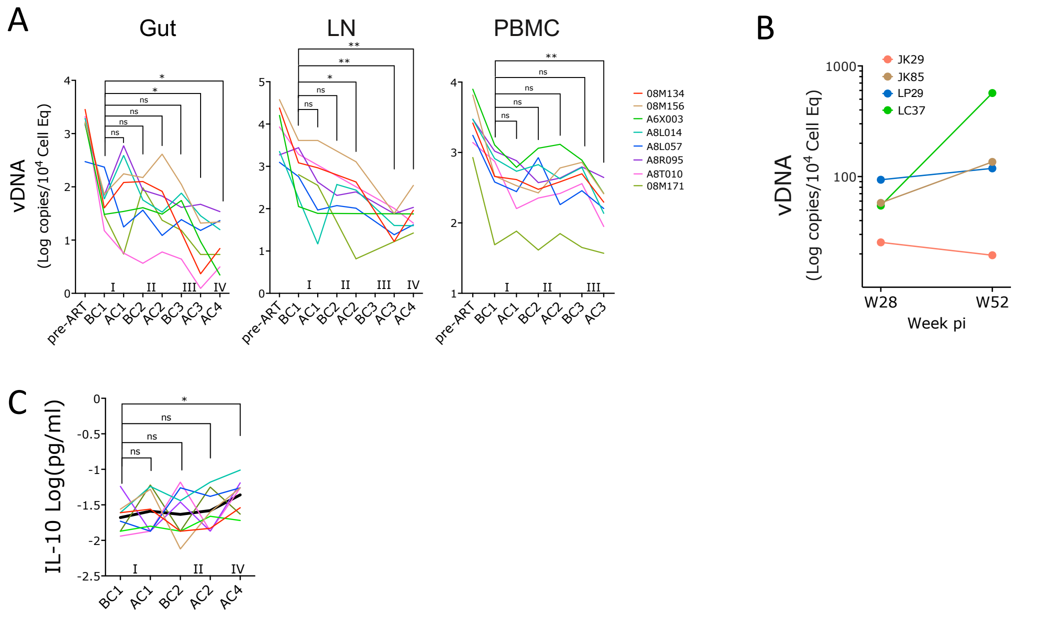


**Figure S5. Galunisertib decreases CA-vDNA with a minor increase in systemic IL-10.** A) Levels of cell-associated (CA)-vDNA per cell equivalent are shown for the pre-ART time point (week 6) and for all the time points before and after the first 3 galunisertib cycles (BC1, AC1, BC2, AC2, BC3, AC3) for PBMC and also for after cycle 4 (AC4) for Gut and LN. pvalues are shown for a mixed effect model and post-hoc Holm-Sidak correction (*p≤0.05 **p≤0.01 ***p≤0.01) BC3 had only 1 value in the LN (08M171) and was not included in the analysis. B) CA-vDNA in PBMC from 4 SIVmac239M2 infected, ART-treated macaques not treated with galunisertib at week 28 vs week 52 post-infection. C) IL-10 levels in plasma at the indicated time point. pvalues are shown for a mixed effect model and post-hoc Holm-Sidak correction (*p≤0.05).

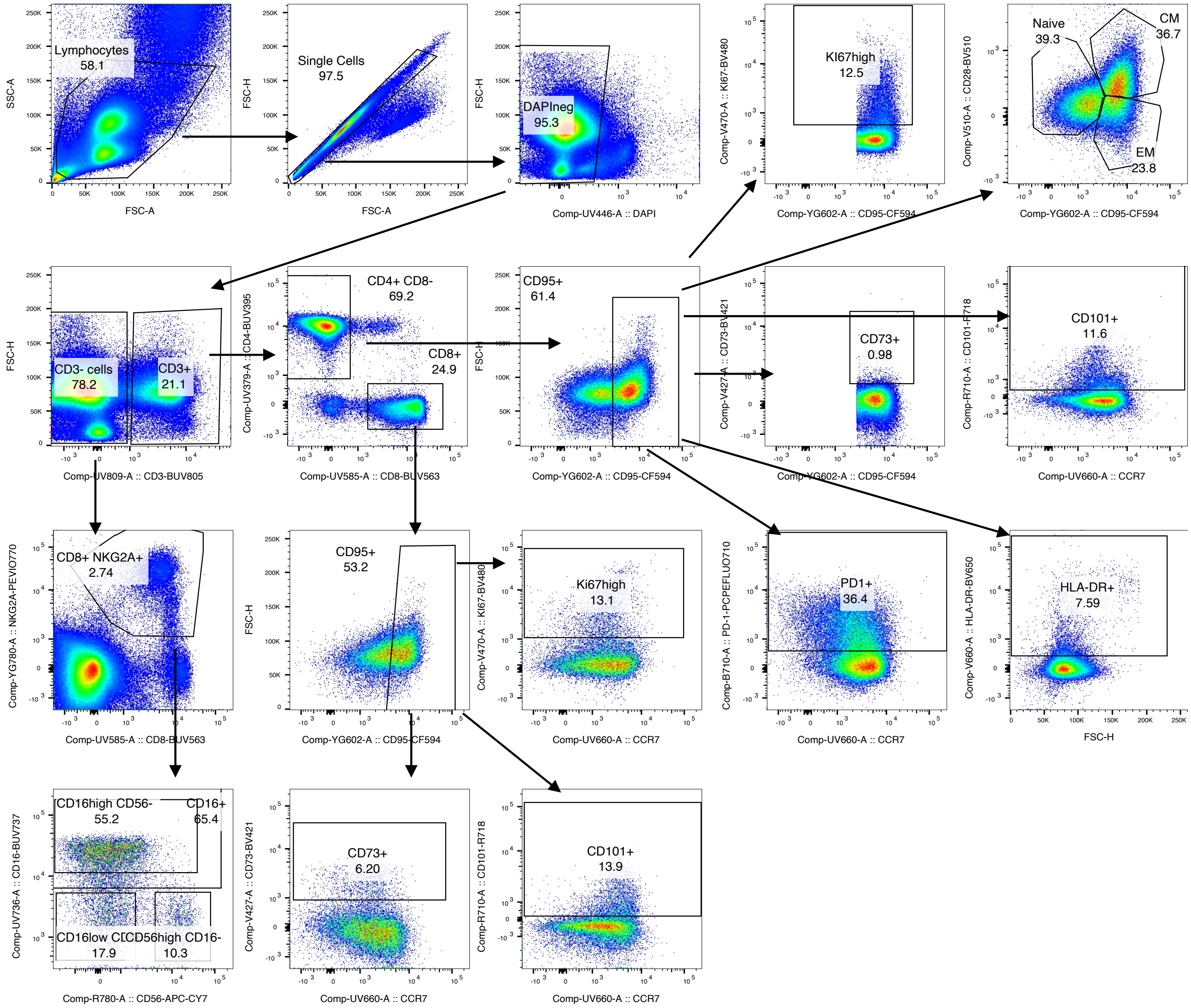


**Figure S6 Gating strategy for classical analysis of 24-color flow.**

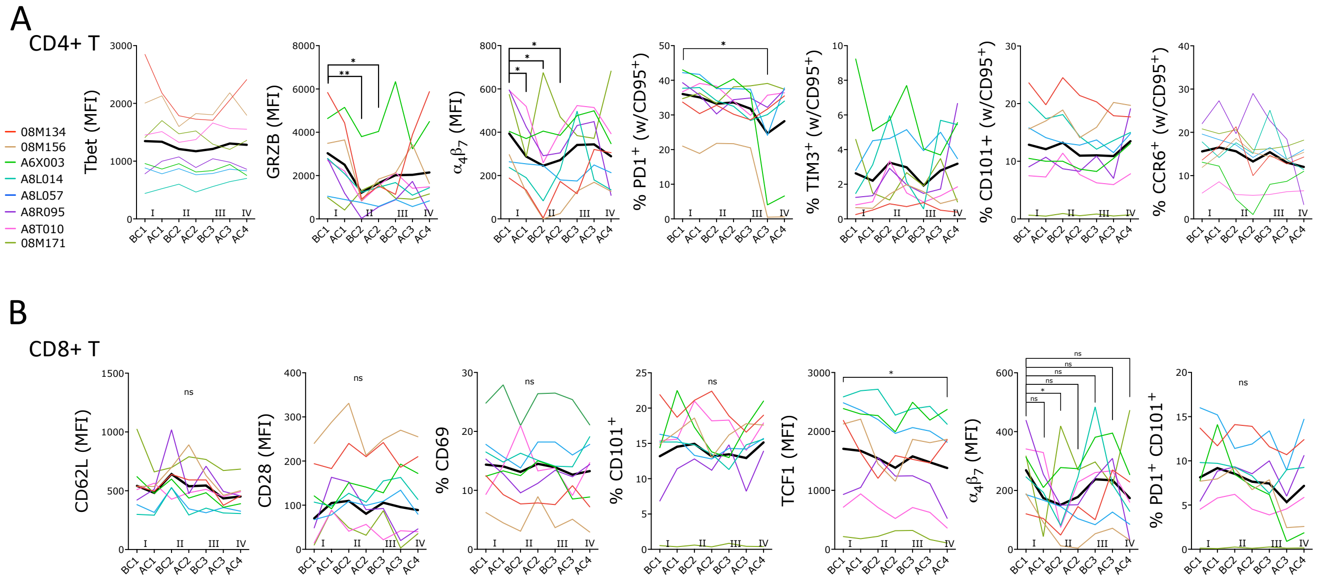


**Figure S7. Decrease in integrin $\alpha 4\beta 7$, but no change in classical activation or exhaustion markers after galunisertib treatment.** A-B) Geometric mean fluorescent intensities (MFI) of each marker and frequency of indicated subset within live, singlets CD3$^+$ CD4$^+$ T cells (A) or within CD3$^+$ CD8$^+$ T cells (B) or CD4$^+$ CD95$^+$ T cells (where indicated on Y label) are shown. Thick black line represents the mean. Changes from baseline (beginning of cycle 1, BC1) are shown only for graphs with at least 1 significant difference (One-way Anova for repeated measures with Holm-Sidak correction for multiple comparisons; *p≤0.05 **p≤0.01 ***p≤0.01). Of note, drop in PD1 expression in cycle 3 for 08M156 and A6X003 is due to in vivo treatment with $\alpha$PD1 Ab. Decline in PD1 is not significant after exclusion of the cycle 3 time point in these 2 macaques.

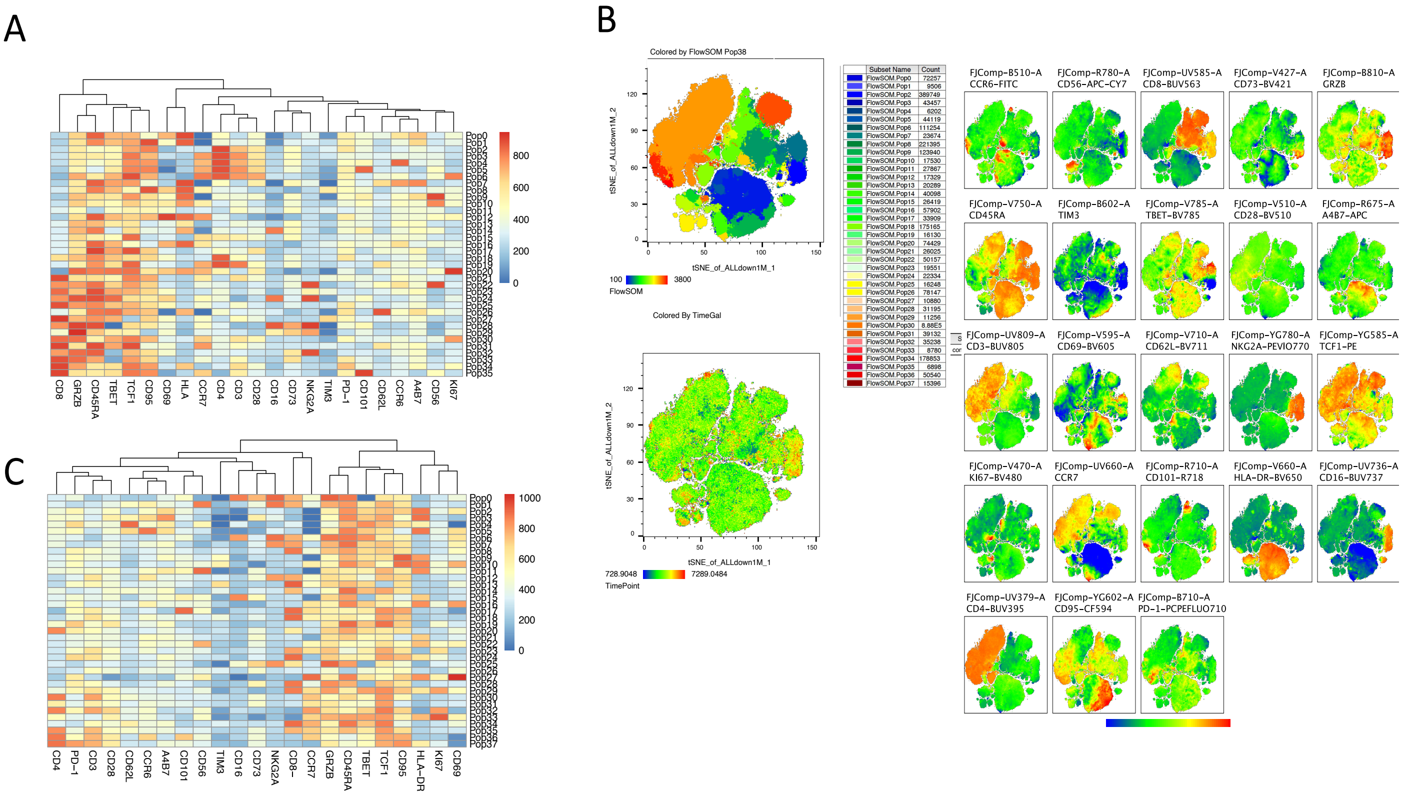


**Figure S8. Heatmaps of populations generated by FlowSOM.** A) FlowSOM was run with phenograph suggested 36 populations on cleaned, live, singlet lymphocytes of BC1 and AC4 data. Heatmap of MFI for each marker in each FlowSOM population is shown. B) tSNE visualization of lymphocyte, live, singlets events after normalization for all time points analyzed (BC1, AC1, BC2, AC2, BC3, AC3 and AC4 for all 8 macaques) with FlowSOM 38 clusters overlaid on the tSNE (top left) or heatmap of each markers MFI (right) or heatmap of time point (blue is BC1 and the progressively more toward red AC4; bottom left) is shown. C) FlowSOM was run with phenograph suggested 38 populations on cleaned, live, singlet lymphocytes of all time points. Heatmap of MFI for each marker in each FlowSOM population is shown.

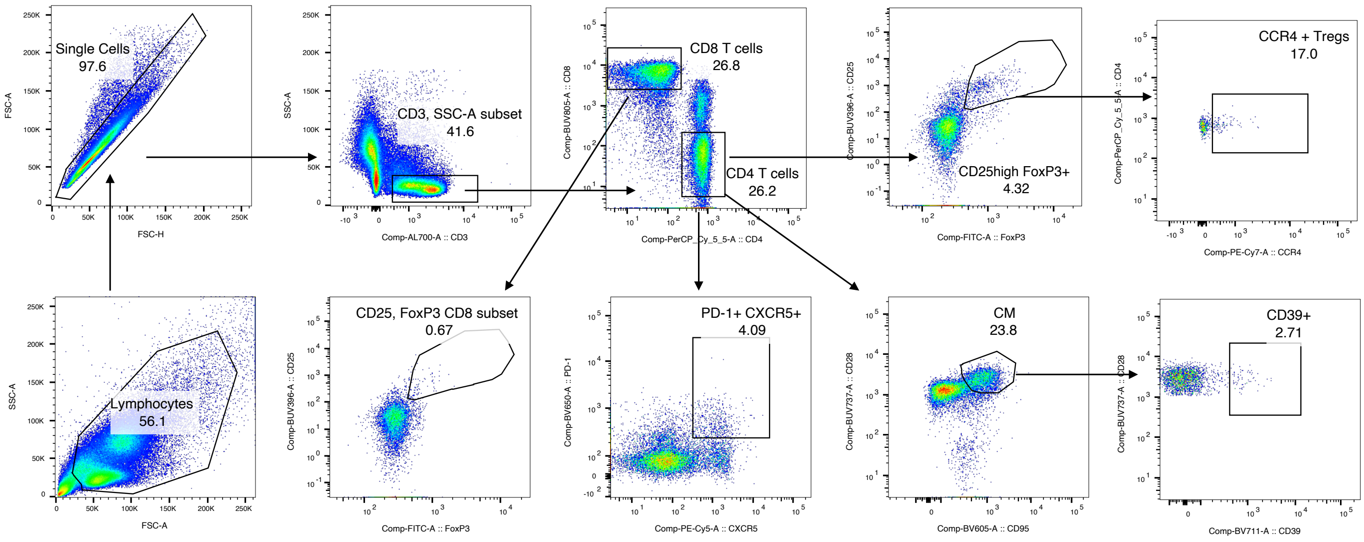


**Figure S9 Gating strategy for analysis of Tfh and Treg frequencies.**

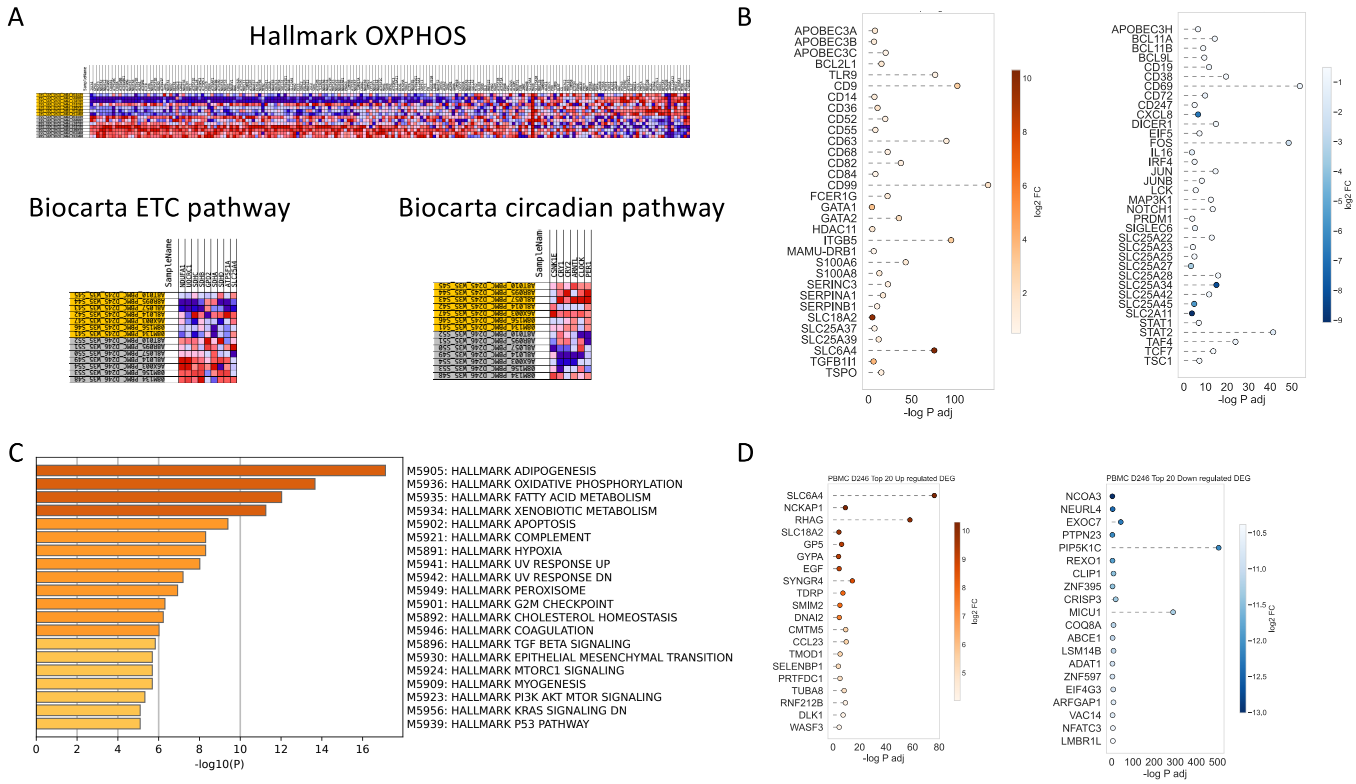


**Figure S10 DEG and pathways enriched 1hr after galunisertib in PBMC.** A) Heatmap of genes in the Hallmark OXPHOS pathways upregulated (red) or downregulated (in each monkey) 1hr after the first dose of galunisetrib. B) DEGs of interest (upregulated left and downregulated right after 1hr). C) Metascape analysis of hallmark gene sets enriched in upregulated DEGs after 1hr of galunisertib. D) List of top 20 DEGs (for log2FC) upregulated or downregulated after 1hr of galunisertib.

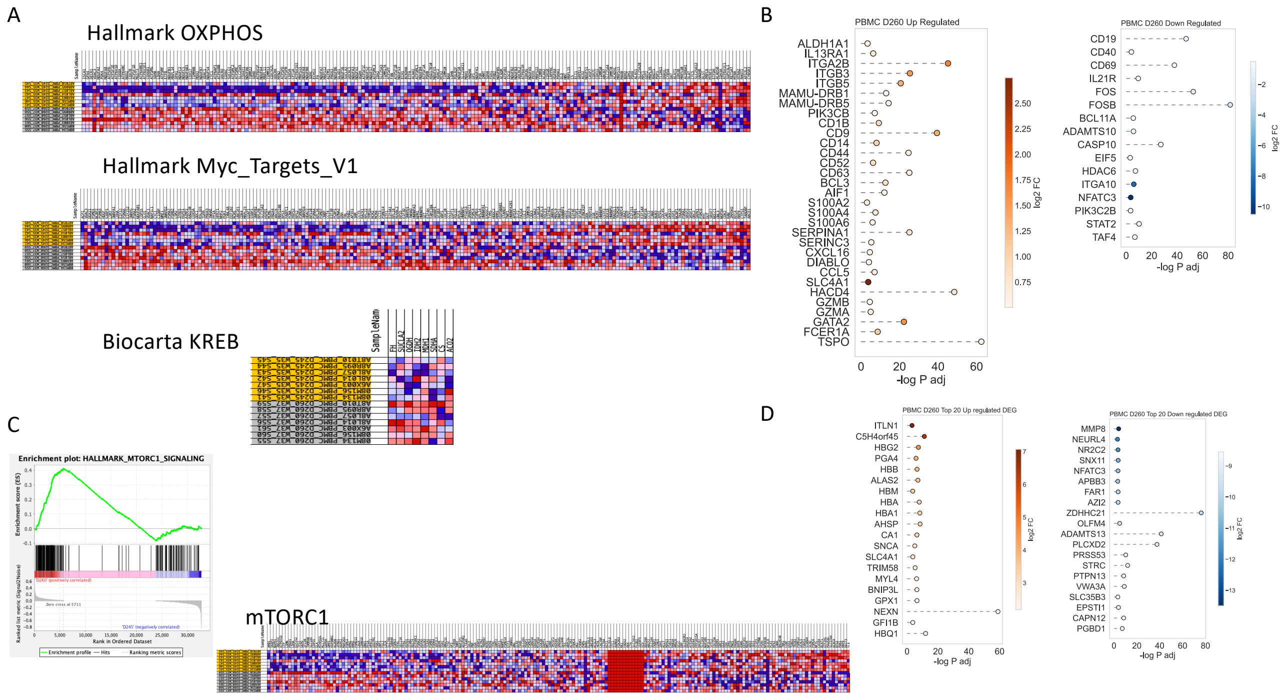


**Figure S11 DEG and pathways enriched after 2 weeks of galunisertib in PBMC.** A) Heatmap of genes in the Hallmark OXPHOS, Myc_Targets_V1 and KREB pathways upregulated (red) or downregulated (in each monkey) after 2 weeks of galunisetrib. B) DEGs of interest (upregulated left and downregulated right after 2 weeks). C) GSEA enrichment plot and heatmap of genes of mTORC1 pathway enriched among hallmark sets. D) List of top 20 DEGs (for log2FC) upregulated or downregulated after 2 weeks of galunisertib.

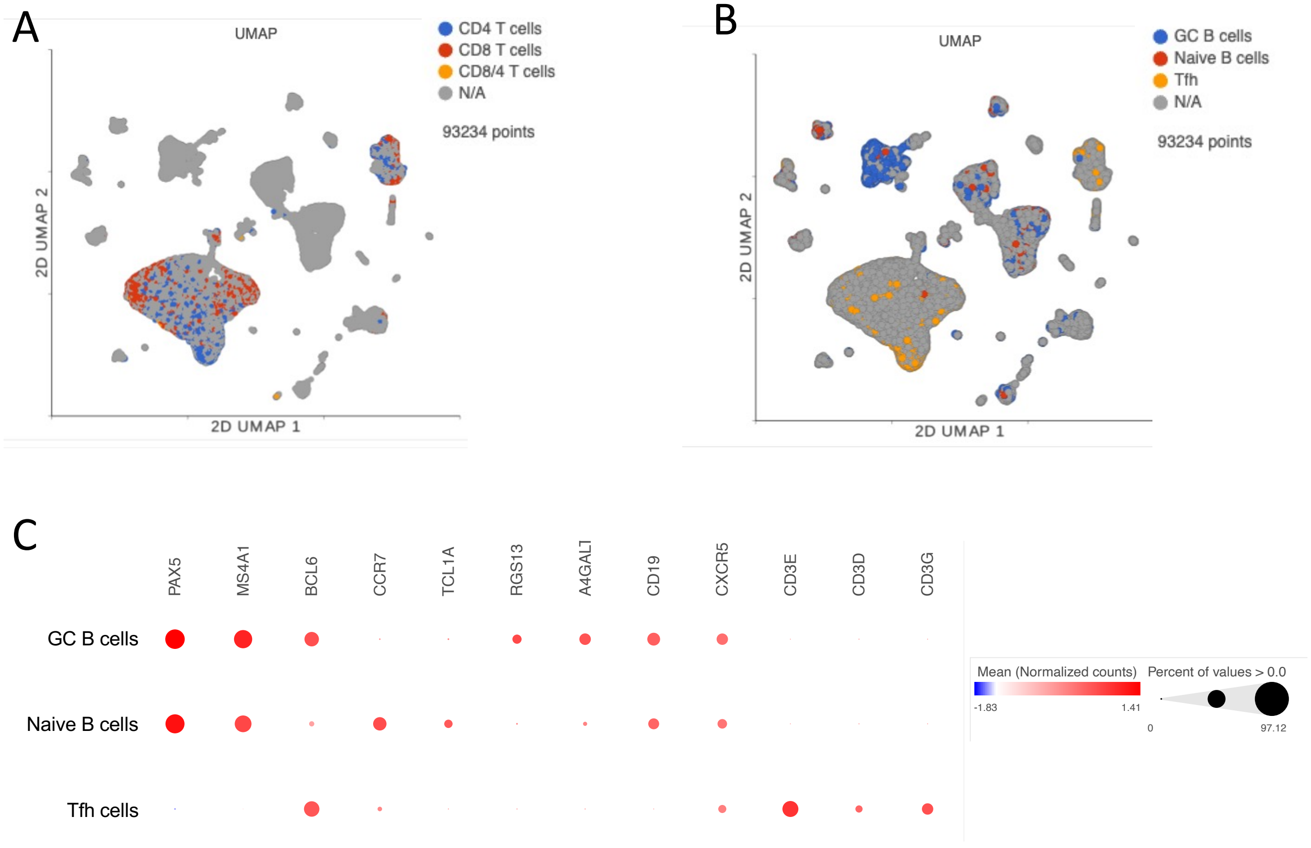


**Figure S12. UMAP and heatmap of cell classification for scRNAseq of lymph node cells before and after galunisertib cycle 1.** A-B**)** UMAP projection of 93234 cells from lymph nodes collected right before and at the end of cycle 1 from all 8 macaques (16 samples). Gene-based classification of CD4 and CD8 T subset (A) and B subsets and Tfh (B) is overlaid on UMAP. In gray are unclassified cells. C) Bubble plot showing expression (mean normalized counts proportional to the color; size proportional to the percentage of cells) of each marker listed in each cell subset. Marker listed are those used for classification of major immune subsets of B cells and Tfh cells.

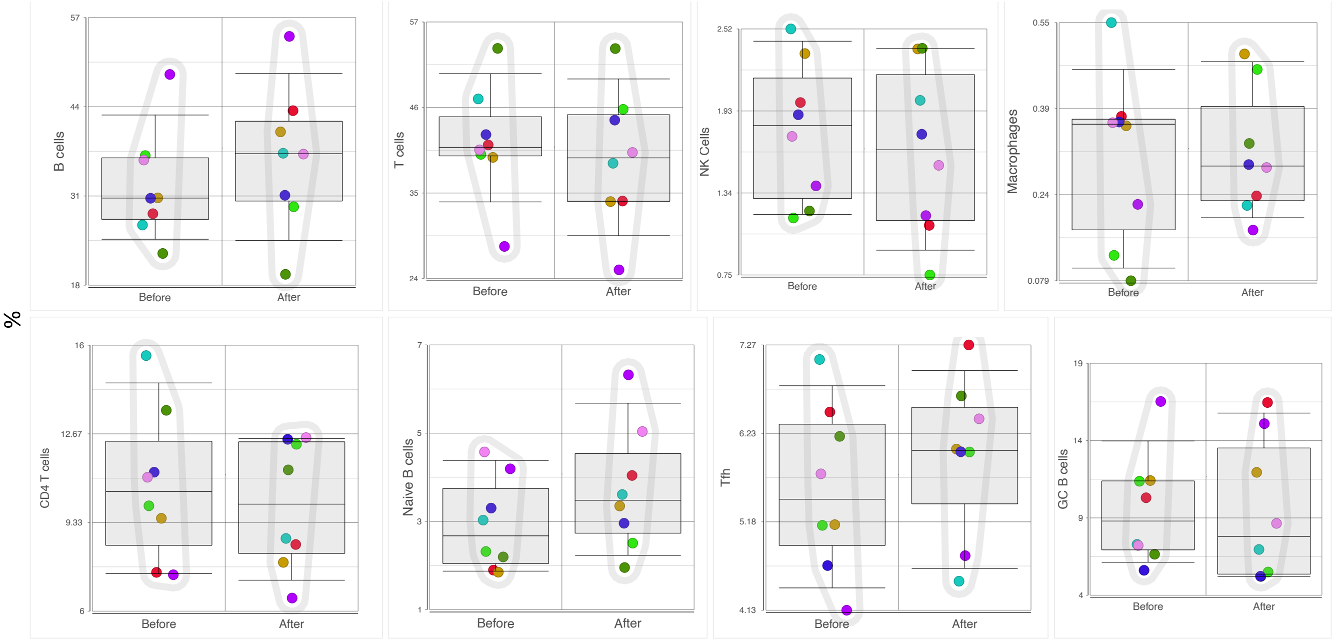


**Figure S13. Frequency of each population within LN before and after cycle 1 with galunisertib.** Cells were classified using gene expression (Fig 6B, 6C and S11C) and the number of cells of a specific type was normalized over the total cell number for each sample. Wilcoxon signed rank test was used to compare frequencies before and after galunisertib (no significant differences p<0.05 were found).

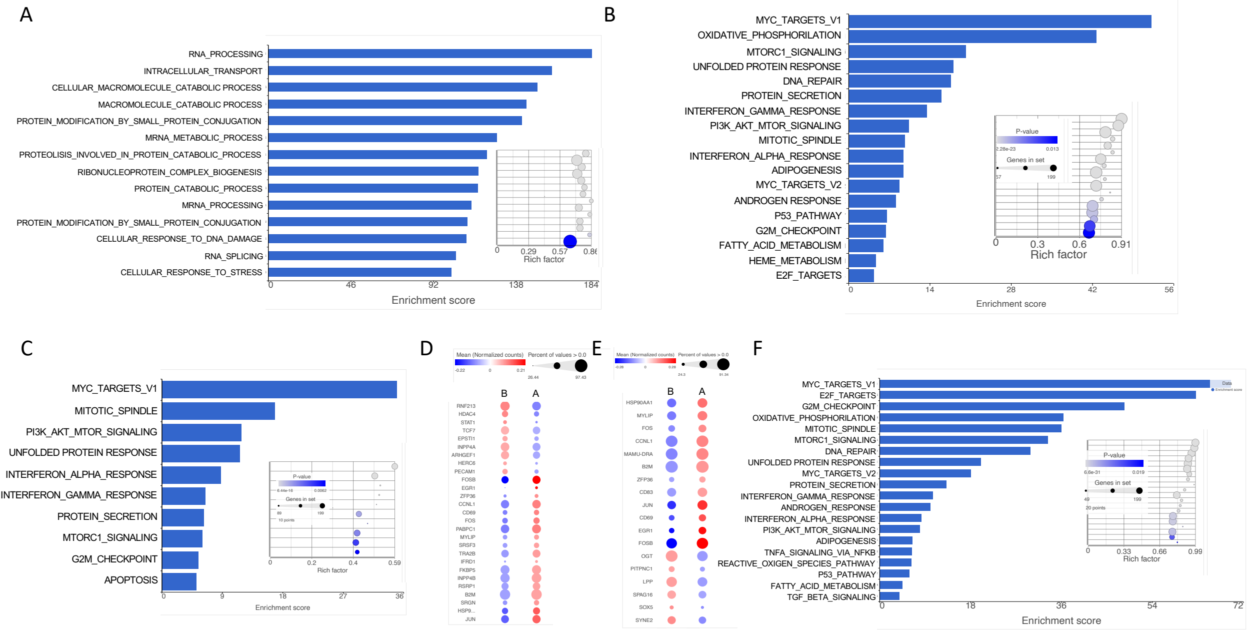


**Figure S14. Metabolic processes are upregulated in both T and B cells after galunisertib in the LN.** A) Pathways enriched in T cells among gene ontology cellular processes collection are listed with ES >100. B) Pathways enriched in CD8 T cells among hallmark sets. C-D) Hallmark pathways enriched (adjp<0.05) and DEGs in Tfh cells (log2fc 0.15). E) DEG in B cells with log2fc 0.2 cutoff. F) Hallmark pathways enriched in B cells. B= before galunisertib, A= After galunisertib

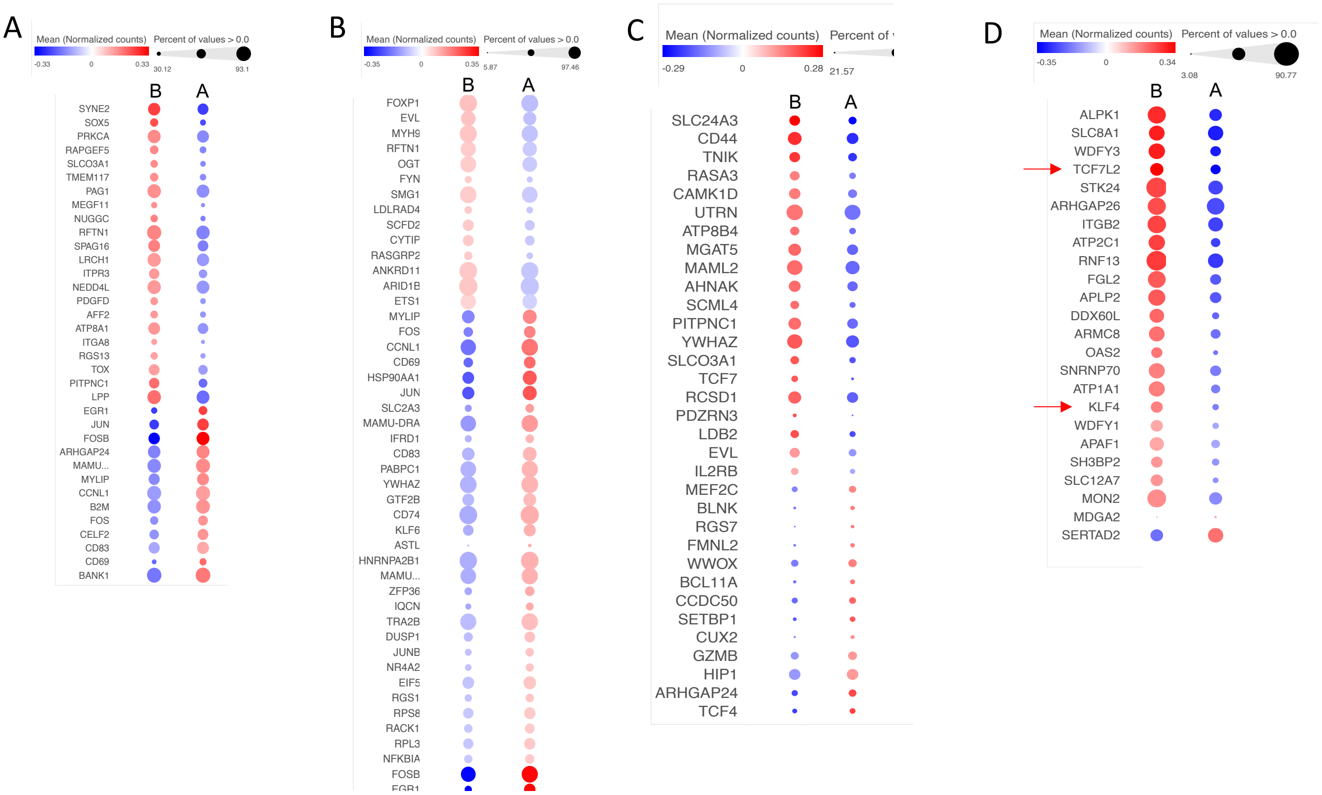


**Figure S15. Galunisertib leads to a switch toward effector/inflammatory phenotype in B cells, NK and macrophages**. A-B) Significant DEGs with a log2FC cut off .15 in GC B cells and naïve B cells. C) Significant DEGs with a log2FC cut off of 0.15 in GC B cells and naïve B cells C) Significant DEG in NK cells with a cutoff log2FC=0.3 D) Significant DEG in Macrophages with a cutoff log2FC= 0.25. B= before galunisertib, A= After galunisertib

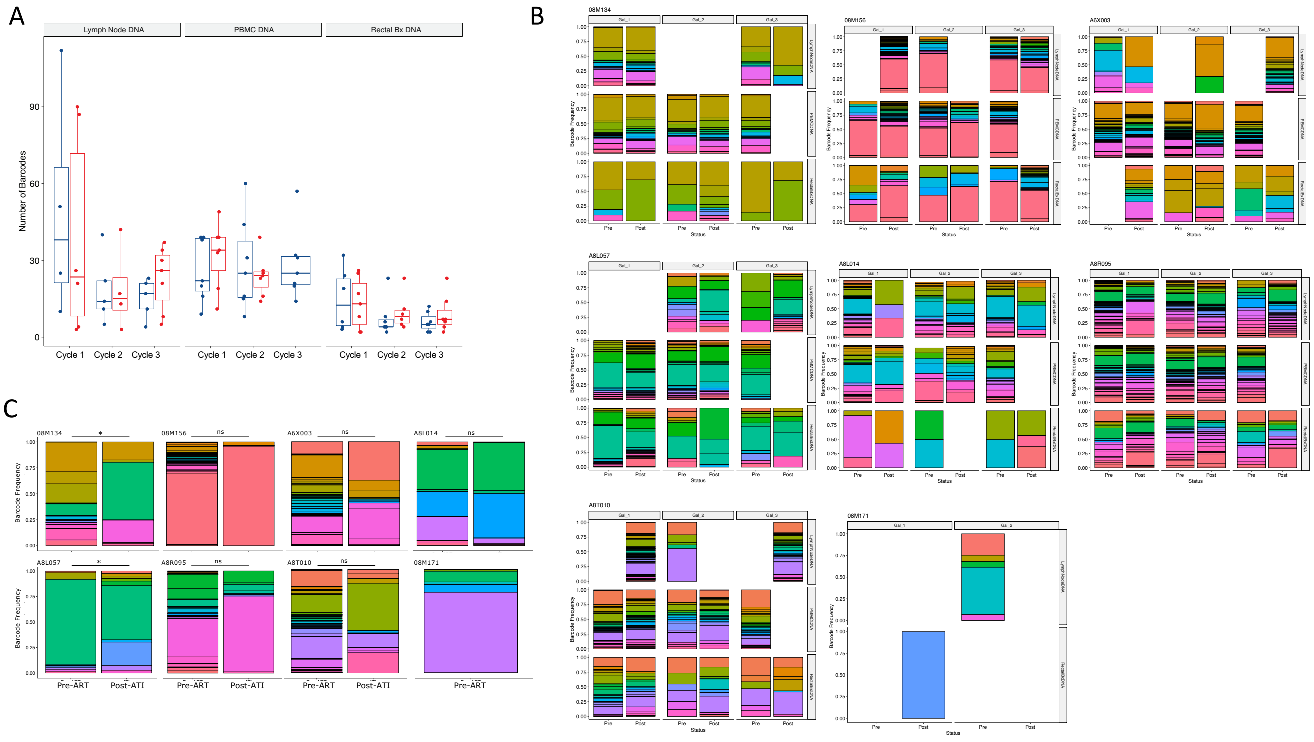

**Figure S16. No changes in barcode numbers with galunisertib, but significant changes in barcode distribution in different tissues**. A) Number of barcodes detected before (blue) and after (red) each galunisertib cycle in the listed tissues. B-C) Barcode frequency distribution in each monkey and tissue in which virus was detected. Each barcode has a different color, and its bar size is proportional to its frequency. Data from before and after each of the first 3 galunisertib cycles (B) or pre-ART vs post-ATI week 6 (C) are shown.

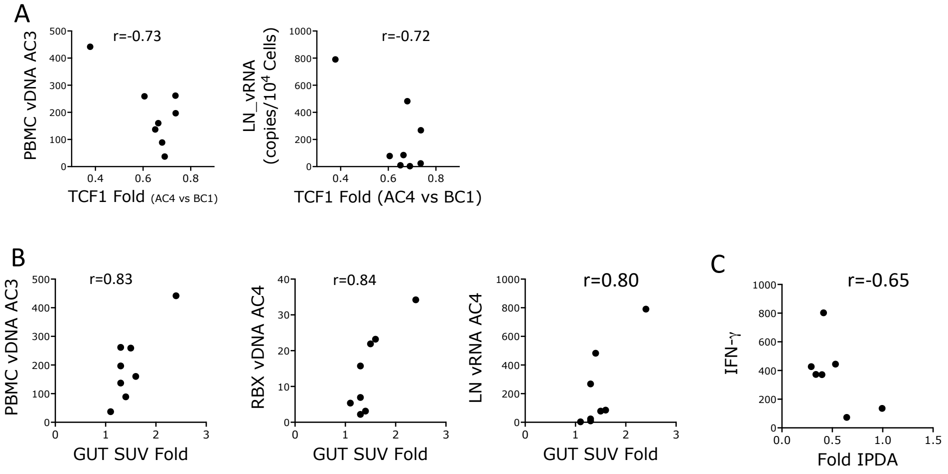


**Figure S17. Correlations between changes in TCF1, Gut SUV and intact pro-virus with virological and immunological parameters.** A) Fold changes in TCF1 MFI are plotted against CA-vDNA copies in PBMC at the end of cylcle 3 and CA-vRNA copies in LN at the end of cycle 4 (AC4). Pearson r is shown B) Fold changes in gut SUV are plotted against CA-vDNA in PBMC at the end of cycle 3 and in rectal biopsies and lymph nodes at the end of cycle 4. Pearson r is shown. A-B) Significant correlations $p<0.05$ C) Fold changes in intact pro-virus are plotted against the avg spot for IFN-$\gamma$ after stimulation with SIV-peptides (env + gag + pol). Pearson r is shown. $p=0.117$

**Supplemental Movie Legends:**

**Supplemental Movies 1-10. Timeline maximum intensity projections (MIP) of fused PET and CT scans are shown for each macaque and to illustrate ROI strategy. Movies 1-8)** Fusions of the whole-body CT and PET scans were generated for each timepoint for each animal. MIPs were generated using the MIM software and set to a numerical scale of 0-1.5 SUVbw visualized with the color scale PET Rainbow. The MIP were scaled to the same physical size using the dimensions to the scanner's visible range, and the scanner bed was removed from the CT scan. The MIP are displayed in chronological order, qualitatively demonstrating the changes in intensity and location of PET signals. Each movie generated in MIM as a series of MIP for one full rotation (~7 seconds) was captured using QuickTime. **Movie 9**) Representative example of MIP serie generated in MIM for full rotation including ROIs as different colored lines for 1 monkey at 1 time point. **Movie 10)** Representative example of MIP serie of Axial viex depicting Ax LN ROI for all scans of a single macaque.

### Supplementary Tables

| Table S1 | Age (yo)* | Sex | Weight * (Kg) | Infection & Route | Treatment | ART-Start Week pi | Cycle 1 Start (BC1) Week pi | Cycle 4 End (AC4) Week pi |
|---|---|---|---|---|---|---|---|---|
| **08M134** | 13 | F | 13.55 | SIVmac239M2 IV | Gal | 6 | 35 | 49 |
| **08M156** | 13 | F | 6.10 | SIVmac239M2 IV | Gal + Nivo | 6 | 35 | 49 |
| **A6X003** | 15 | F | 7.85 | SIVmac239M2 IV | Gal + Nivo | 6 | 35 | 49 |
| **A8L014** | 13 | F | 7.70 | SIVmac239M2 IV | Gal | 6 | 35 | 49 |
| **A8L057** | 13 | F | 9.50 | SIVmac239M2 IV | Gal | 6 | 35 | 49 |
| **A8R095** | 13 | F | 8.20 | SIVmac239M2 IV | Gal | 6 | 35 | 49 |
| **A8T010** | 13 | F | 5.05 | SIVmac239M2 IV | Gal | 6 | 35 | 49 |
| **08M171** | 13 | F | 7.35 | SIVmac239M2 IV | Gal | 6 | 32 | 46 |
| **Average** | **13.2** | | **8.12** | | | | | |

*At beginning of the study

**Table S1. List of macaques used for the main study described in the manuscript**

**Table S2. Hematology Data – See excel file**

**Table S3 Chemistry Data – See excel file.**

| PBMC High-Dim | Marker | Clone | Maker |
|---|---|---|---|
| **DAPI** | FVS440UV | Live Dead | BD |
| **BUV395** | CD4 | L200 | BD |
| **BUV563** | CD8 | RPA-T8 | BD |
| **BUV661** | CCR7 | 3D12 | BD |
| **BUV737** | CD16 | 3G8 | BD |
| **BUV805** | CD3 | SP34-2 | BD |
| **BV480** | Ki67** | B56 | BD |
| **BV510** | CD28 | 28.2 | Biolegend |
| **BV605** | CD69 | FN50 | Biolegend |
| **BV650** | HLA-DR | L243 | BD |
| **BV711** | CD62L | SK11 | BD |
| **BV750** | CD45RA | 5H9 | BD |
| **BV786** | Tbet | 4B10 | Biolegend |
| **AF488** | CCR6 | G034E3 | Biolegend |
| **BB630** | Tim-3 | 7D3 | BD |
| **BB700/PCPeFluo710** | PD-1 | eBioJ105 | eBioscience |
| **BB790** | GRZB** | GB11 | BD |
| **PE** | TCF-1** | S33-966 | BD |
| **PE-CF594** | CD95 | DX2 | Biolegend |
| **PE-Vio770** | NKG2A | REA110 | Mylteny |
| **APC** | a4b7 | Act-1 | NHP |
| **AF700/R718** | CD101 | V7.1 | BD |
| **APC-Fire750** | CD56 | HCD56 | Biolegend |

| Treg/Tfh Panel | | | |
|---|---|---|---|
| **FITC** | FOXP3 | PCH101 | EBioscience |
| **PCP-Cy5.5** | CD4 | L200 | BD |
| **BV421** | CTLA4 | BNI3 | Biolegend |
| **BV510** | L/D | N/A | BD |
| **BV605** | CD95 | DX2 | Biolegend |
| **BV650** | PD-1 | EH12.2H7 | Biolegend |
| **BV711** | CD39 | A1 | Biolegend |
| **BUV395** | CD25 | BC96 | BD |
| **BUV737** | CD28 | CD28.2 | BD |
| **BUV805** | CD8 | RPA-T8 | Biolegend |
| **PE-Cy5** | CXCR5 | C398.4A | Biolegend |
| **PE-Cy7** | CCR4 | L291H4 | Biolegend |
| **APC** | TCF-1 | s33-966(RUO) | BD |
| **AL700** | CD3 | SP34-2 | BD |

**Table S4 List of Abs used for phenotyping.**

**Table S5 Raw data from flow cytometry classical analysis and MFIs.** See Table S5 excel file

| RI=Right inguinal<br>LI= Left inguinal | | | Cycle 1 | | Cycle 2 | | Cycle 3 | | Cycle 4 |
|---|---|---|---|---|---|---|---|---|---|
| LA= Left Axillary | Peak VL | pre-ART | BC1 | AC1 | BC2 | AC2 | BC3 | AC3 | AC4 |
| RA= Right Axillary | W2 | W6 | W35 | W37 | W39 | W41 | W43 | W45 | W 49 |
| 08M134 | RI,RA | RA | RA | RI | LA | LA | RA | RA | RA/LA |
| 08M156 | RI,RA | LI | RA | LI | LA | LA | RA | RA | RA/LA |
| A6X003 | RI | RA | RA | RI | LA | LA | RA | LA | RA/LA |
| A8L057 | RA,LA | RA | RA | RI | LA | LA | LA | LA | RA/LA |
| A8L014 | LI,RI | RI | RA | RI | LA | LA | LA | LA | RA/LA |
| A8R095 | LI,RI | RI | RA | LI | LA | LA | RA | RA | RA/LA |
| A8T010 | RA,LA | RA | RA | LA | LA | RA | RA | LA | RA/LA |
| O8M171 | | | LA | LA | RA/RI | LA | LA | RI | RA/LA |

**Table S6 List of**
